# Genomic-Based Prediction of Exopolysaccharide Composition and Structure: Insights from *Rhizobium* and *Sinorhizobium* Species

**DOI:** 10.64898/2026.08.21.746188

**Authors:** Joris Tulumello, Justine Long, Wafa Achouak, Marie-Line Garron, Nicolas Terrapon, Thierry Heulin

## Abstract

Bacterial exopolysaccharides (EPS) are key components in biofilm formation, stress protection, and symbiosis in Rhizobiaceae. While EPS structural diversity is extensive, experimental characterization remains limited. In this study, we experimentally determined and compared four distinct EPS structures produced by ten *Rhizobium alamii* strains. Using genomic data, we bioinformatically identified supra-operonic clusters (SOCs) responsible for these EPS biosynthesis. We introduced a computational framework to predict, score, and compare EPS SOCs across 84 *Rhizobium* and *Sinorhizobium* species, linking gene content to structural and functional EPS diversity. A total of 743 EPS SOCs was selected for network analyses, allowing the identification of 36 major groups of orthologous EPS SOCs, successfully recovering all known EPS biosynthetic loci and two novels SOCs potentially encoding uncharacterized EPS (xEPS-I, xEPS-II). Profiles of EPS SOCs correlated with taxonomical groups, with a single EPS SOC conserved through all 84 genomes and distinct additional EPS SOCs depending on the group, but do not strictly explain symbiotic capacity. Genetic comparisons of transporters (Wzx, Wzy) and glycosyltransferase sequences indicated these proteins as key markers of EPS structure. Overall, this computational framework accurately identified and classified EPS SOCs, providing a scalable, genome-based method for predicting EPS biosynthetic potential in *Rhizobiaceae* and usable in other microbial genera.

**Author Summary:** We developed a computational approach to predict how beneficial soil bacteria produce natural exopolysaccharides (EPS) that help them survive and interact with plants. These macromolecules, which form protective coatings and enable partnerships with crops, are currently difficult and expensive to study using traditional laboratory methods. By examining the genetic blueprints of 84 bacterial species from the Rhizobiaceae, we identified the key gene clusters responsible for producing these EPS, including two that were previously unknown. Our findings show that while some genetic patterns are widely shared among many bacteria, others are unique to specific types, shaping their individual abilities and characteristics. This research provides a faster, more cost-effective way to explore the vast diversity of these important macromolecules across different bacterial species. These insights could enable enhanced bacterial treatments that boost crop growth and resilience, particularly against water stress. Additionally, our method offers a valuable template for studying similar processes in other beneficial microorganisms, ultimately advancing our understanding of how they contribute to plant health, soil fertility, and the development of more sustainable farming practices worldwide.

## Introduction

Bacterial exopolysaccharides (EPS) are essential extracellular biopolymers facilitating interactions with their environment. They play critical roles in biofilm formation, protection against chemical or physical stress, and the retention of water and nutrients. In members of the Rhizobiaceae family, EPS are crucial for symbiosis establishment and bacterial survival throughout symbiotic stages [1]. Notably, some free-living and EPS-producing Rhizobiaceae do not establish symbiotic relationships but promote plant growth, possibly via EPS and other mechanisms such as phytohormone action, or modulation of root-associated microbiota [2–4].

A single bacterial strain can produce diverse EPS with distinct structures and functions. For instance, *Rhizobium leguminosarum* bv. *viciae* 3841 (newly renamed *Rhizobium johnstonii* 3841) synthesizes four distinct EPS: one homopolysaccharide (a β-glucan like cellulose), and three heteropolysaccharides (a glucomannan, an acidic EPS, and an EPS of unknown structure) [5]. Mutational studies indicate the glucomannan may bind with root hair lectins, the cellulose supports bacterial aggregation on root hairs, and the acidic EPS is essential to symbiosis establishment [5]. EPS are composed of monosaccharide-repeating units, often decorated by non-carbohydrate substituents, showing extensive variability between species and even strains. For example, *Rhizobium alamii* strains can produce four distinct EPS types (**Figure 1A**) [6–8]. Similarly, *R. leguminosarum* bv. *viciae* strains 3841 (Rlv3841) and 248 (Rlv248) synthesize different EPS structures, with an identical core chain but variations in the lateral branch (**Figure 1B**) [9,10]. Remote species can however produce EPS with highly similar structures, as *Rhizobium etli* CFN42 and *R. leguminosarum* bv. *trifolii* TA1 share Rlv3841 EPS structure. Similarly, *Sinorhizobium meliloti* 1021 (Sme1021) produces two EPS, a succinoglycan (EPS-I) and a galactoglucan (EPS-II), while *Rhizobium favelukesii* LPU83 produces the same EPS-I, and *Sinorhizobium fredii* HH103 an EPS with notable similarities (**Figure 1B**) [11–13].

**Figure 1:**
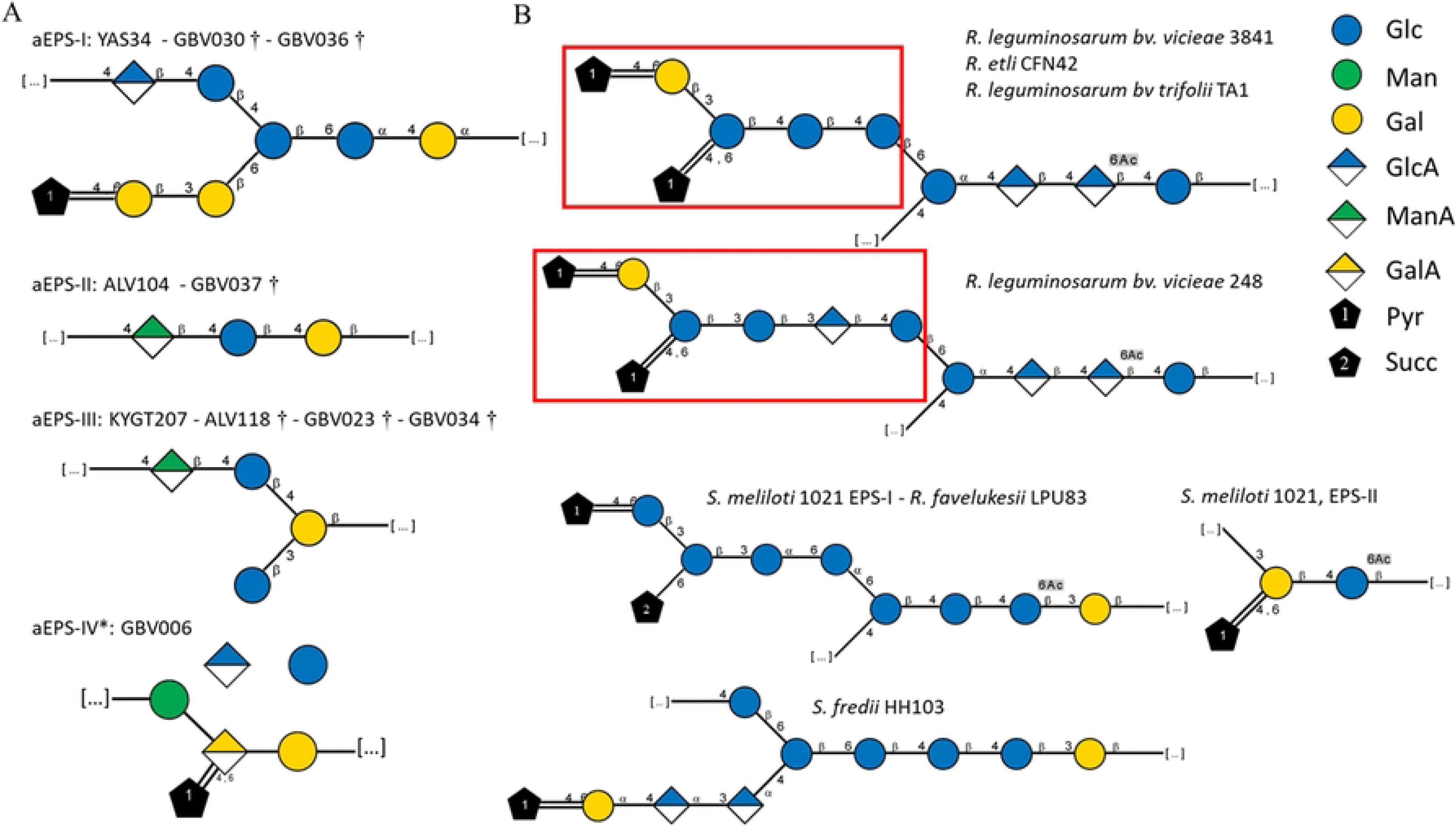
EPS structures presented in Symbol Nomenclature for Glycans (SNFG) format. EPS from (A) *R. alamii* strains analyzed previously and, in this study, (denoted by †), and (B) *Rhizobium* and *Sinorhizobium* strains. For *R. alamii*, different strains listed above a single EPS structure produce the same EPS. The structure of aEPS-IV (indicated by *) is only partially characterized; the precise positions of Glc and GlcA, as well as osidic bonds, remain undetermined. The position of acetate groups on *R. alamii* EPS structures could not be definitively assigned. Brackets ‘[…]’ at both extremities of a repeating unit indicate that another repeating unit is linked via an osidic bond between these carbohydrate residues, forming the EPS chain. Carbohydrates enclosed within the brackets constitute the core structure of the repeating unit, while carbohydrates outside the brackets form the lateral branch. The core structure of the repeating units in Rlv3841 and Rlv248 are identical, whereas their lateral branches (highlighted by red squares) differ.

Determining an EPS structure requires extraction, purification, and chemical analyses such as degradation reactions and/or NMR spectroscopy in D_2_O, which are time-consuming and costly. Consequently, EPS structures are not systematically characterized for all bacterial strains. In 2020, Knirel and Van Calsteren compiled 29 different polysaccharide structures representing EPS or capsular polysaccharide (CPS) from 19 *Rhizobium* and 10 *Sinorhizobium* strains [14]. At that time, only four of these strains had complete genomes publicly available. However, as of January 2026, the NCBI genome database lists 2,263 *Rhizobium* and 915 *Sinorhizobium* genomes. The rapidly decreasing cost of genome sequencing should facilitate genome-based studies of EPS biosynthesis, enabling predictive modeling of EPS production and expanding the exploration of EPS diversity, while helping to better target labor-intensive chemical characterization approaches.

In heteropolysaccharide biosynthesis and export, the repeating oligosaccharide unit is assembled on a lipid carrier through the sequential action of specific glycosyltransferases (GTs). These enzymes utilize activated monosaccharide substrates, specifically nucleotide-monosaccharides, such as uridine diphosphate-glucose (UDP-Glc), UDP-galactose (UDP-Gal), UDP-galacturonic acid (UDP-GalA), or guanosine diphosphate-mannose (GDP-Man) [15]. The enzymes responsible for synthesizing these nucleotide-monosaccharide substrates are collectively referred to as precursor enzymes (**Figure 2A**). The KEGG pathways, <u>map520</u> and <u>map541</u> [16], provide a comprehensive overview of the biosynthetic routes for the majority of nucleotide-monosaccharides essential for EPS biosynthesis. The biosynthesis of nucleotide-activated monosaccharides primarily originates from two central metabolic intermediates: glucose-1-phosphate (Glc-1P) and fructose-6-phosphate (Fru-6P). Glc-1P serves as the precursor for the formation of UDP-Glc, which is subsequently epimerized to UDP-Gal by ExoB in Sme1021 [17]. Additionally, UDP-Glc is converted to UDP-GlcA, through the action of RkpK, followed by its transformation into UDP-GalA, a reaction catalyzed by IspL in *S. meliloti* Rm41 [18]. Additionally, Glc-1P is directed toward the synthesis of TDP-rhamnose (TDP-Rha) via the conserved *rmlABCD* operon, initially characterized in *Salmonella enterica* [19]. Fru-6P acts as the metabolic precursor for distinct pathways involved in the biosynthesis of GDP-activated monosaccharides. In *S. fredii*, the sequential enzymatic activity of NoeJ and NoeK convert Fru-6P into GDP-Man, which is further modified to GDP-fucose by NoeL [20]. Alternatively, in *Sphingomonas elodea* ATCC 31461, a homologous RkpK-like enzyme catalyzes the formation of GDP-mannuronic acid (GDP-ManA) from Fru-6P [21].

**Figure 2:**
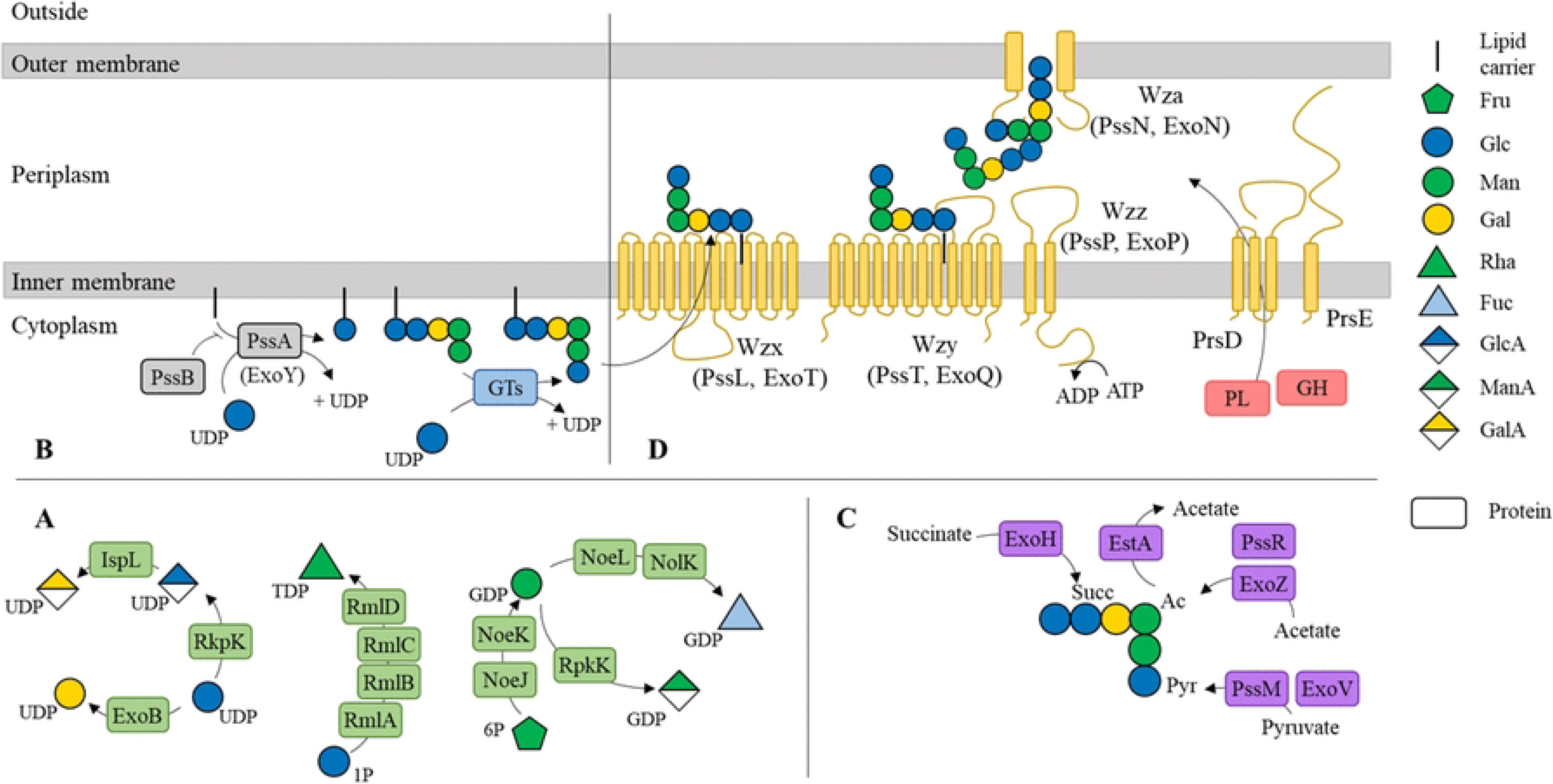
Wzx/Wzy-dependent model EPS synthesis. (**A**) The process begins by the production of nucleotide-activated carbohydrate precursors by enzymes (in green), followed by (**B**) the assembly of the repeated unit, initiated by priming enzymes (in grey) that attach the first carbohydrate to a lipid carrier, with subsequent carbohydrates added by glycosyltransferases (GTs, in blue). (**C**) The repeated unit can be further modified with enzymes (in purple) that add decorations such as acetate, pyruvate, or succinate groups. (**D**) Export of the completed repeated unit is mediated by proteins (in yellow), including Wzx, which transports the lipid-linked repeated unit across the inner membrane; Wzz, which recruits Wzy for polymerization; and Wza, which facilitates export through the outer membrane. A glycoside hydrolase (GH) or polysaccharide-lyase (PL) (in pink) can be transported to the periplasm via the PrsDE proteins, to cleave the glycosidic bonds and modulate EPS length. Only PssLPNT proteins are part of the canonical Wzx/Wzy model, while PrsDE represents an incomplete type 1 secretion system. Proteins labelled ‘Pss’ are derived from Rlv3841, and those labelled with ‘Exo’ are from Sme1021 EPS SOC.

The assembly of heteropolysaccharides is initiated by the priming proteins PssA and PssB. PssA is responsible for attaching the first monosaccharide to the lipid carrier, while PssB regulates the subsequent assembly of the repeating units (**Figure 2B**) [10]. The elongation of the EPS chain is then catalyzed by distinct GTs, which vary depending on the bacterial species (**Figure 2B**). GTs are classified into families based on amino acid sequence similarity in the Carbohydrate-Active enZymes database, CAZy [22]. While many GT families exhibit high substrate specificity, - e.g., GT3 transferring glucose (UDP-Glc) during glycogen synthesis-the largest GT families, GT2 and GT4, show much broader substrate and functional diversity, thereby limiting the predictive power of biosynthetic pathway annotation. Nevertheless, these families share conserved catalytic mechanism: GT2 enzymes catalyze inverting reactions producing β-bonds from α-activated monosaccharide, whereas GT4 enzymes catalyze retaining reactions that preserve the α-conformation of the activated donors [15].

The EPS repeating units can further be decorated with substituents such as acetate, pyruvate, or succinate, through the action of specialized enzymes (**Figure 2C**). Acetyltransferases catalyze the incorporation of acetate groups, ketal-pyruvate-transferases add pyruvate and succinyltransferases introduce succinate [23,24]. These modifications generally occur prior to polysaccharide polymerization and export [25].

The export of EPS via the Wzx/Wzy-dependent model involves four key protein families: Wzx flippases, Wzy polymerases, Wzz polysaccharide co-polymerases, and Wza outer membrane transporters [26] (**Figure 2D**). In this model, Wzx flippases translocate the lipid-linked repeating units across the inner membrane, while Wzy polymerases elongate the EPS chain by connecting these repeating units, Wzz co-polymerases regulate the chain length by recruiting Wzy through ATPase activity, and Wza transporters form an outer membrane channel to facilitate EPS export (**Figure 2D**). Additionally, other CAZymes may participate in EPS biosynthesis, such as glycoside hydrolase (GH) and polysaccharide lyase (PL). These enzymes are transported to the periplasm via the PrsD (ABC transporter) and PrsE (type-I secretion system adaptor), where they cleave glycosidic linkages to produce low molecular weight EPS from larger polymers, such as PssW (GH10) in Rlv3841 and ExoK (GH16_21) in Sme1021 [27]. The coordinated action of these enzymes is essential for efficient EPS production.

Genes encoding the biosynthetic enzymes and transport machinery involved in EPS production are often organized into multiple co-transcribed operons, referred to as supra-operonic clusters (SOCs) [28], enabling coordinated regulation and synchronized EPS biosynthesis. In the Rlv3841 genome, most *pss* genes are clustered within a 9.8 kb region, hereafter designated rEPS-I, whereas the priming glycosyltransferases *pssA* and *pssB* are located approximately 90 kb away, separated by around 80 genes [10]. In Sme1021, two SOCs located on the symbiotic megaplasmid pSymB, are hereafter designated sEPS-I (the *exs*/*exo* region) and sEPS-II (the *wgx/exp* region) [29]. The rEPS-I SOC lacks genes involved in precursor synthesis and priming reactions but encodes nine GTs (*pssCDEFGHIJS*), three decorating enzymes (*pssKMR*), the four Wzx/Wzy transporters (*pssLPNT*), one GH (*pssW*), one polysaccharide lyase (*plyA*), and two auxiliary transporters (*prsDE*) [10]. The sEPS-I SOC contains two precursor synthesis genes (*exoBN*), the priming GT *exoY* (homologous to *pssA*), six GTs (*exoALMOUW*), three decorators (*exoHVZ*), the four Wzx/Wzy transport proteins (*exoTPFQ*), one GH (*exoK*), and two auxiliary secretion proteins (*exsAB*) [23,24]. In contrast, the sEPS-II SOC encodes four precursor synthesis proteins (*expA7, exp8, exp9, exp10*), five GTs (*expA23, expC, expE2, expE4, expE7*), three decorators (*expA4, expA5, expE3*), and two transport proteins (*expD1, expD2*) constituting a Type 1 secretion system based on ABC transporter model [30].

Historically, the identification of SOCs involved in EPS synthesis in rhizobia relied on mutagenesis approaches coupled with the screening of EPS-deficient mutants. For instance, Tn5 mutagenesis in *R. alamii* YAS34 identified the gene EU184019, encoding a GT4, as essential for EPS production [31]. Currently, EPS-associated SOCs (EPS-SOCs) in newly sequenced strains have been mainly identified through comparative genomics using BLAST-based alignments of either individual genes or entire SOC regions. Although effective for detecting homologs of previously characterized SOCs, this strategy remains poorly suited for the discovery of novel EPS biosynthetic clusters [32]. Comparative analyses have revealed a generally conserved EPS-SOC architectures among *R. leguminosarum* strains, albeit with variations in GT content and the occasional presence of additional copies of *pssT* [10]. Horizontal gene transfer from *S. meliloti* has also been proposed to explain the presence of two sEPS-I biosynthesis SOCs in *R. favelukesii*, whose involvement in EPS production was subsequently confirmed by targeted gene deletion [11].

This study aimed to develop a comprehensive framework to: (i) automate the prediction of EPS-SOC directly from genomic data, (ii) implement a scoring system to evaluate EPS biosynthetic potential, and (iii) perform comparative analyses of predicted EPS-SOCs across diverse *Rhizobium* and *Sinorhizobium* strains. To this end, a training dataset comprising 10 *R. alamii* strains with four distinct EPS structures, together with the reference genomes Rlv3841 and Sme1021 was used to calibrate the approach. The resulting predictor was applied to 87 bacterial genomes to explore relationship between SOC gene content and detailed EPS structural features.

## Results and Discussion

### Composition and structure of exopolysaccharides in *Rhizobium alamii*

Obtaining the complete structural characterization of the EPS produced by a given *R. alamii* strain, hereafter referred to as aEPS, requires extensive and time-consuming experimental procedures. These include production in a medium containing 20 g/L glucose, purification by centrifugation to remove bacterial cells, precipitation with ethanol, and finally lyophilization before NMR analysis. Three complete EPS structures have previously been resolved for strains YAS34 (aEPS-I), ALV104 (aEPS-II), and KYGT207 (aEPS-III) [6–8] (**Figure 1A**). The full structure of the EPS produced by strain GBV006, aEPS-IV full structure has not yet been elucidated, and only compositional data with partial structural information are currently available. Using HPLC-MS, we analyzed the *R. alamii* EPS from additional strains GBV023, GBV030, GBV034, GBV036 and GBV037, as well as from formerly reported ALV104 (**Table S1**). These analyses revealed four distinct EPS groups. Strains ALV104 and GBV037 likely produced aEPS-II, characterized by an average repeating unit of 1.0 Glc, 0.97 galactose (Gal) and 0.72 mannuronic acid (ManA) (**Table S1**). Since the known ALV104 EPS structure contains one ManA per repeating unit (**Figure 1A**) [6], these results suggest that HPLC-MS analysis underestimates ManA content. Similarly, strains GBV023 and GBV034 likely synthesized aEPS-III, with average repeating unit of 2.0 Glc, 1.05 Gal and 0.75 ManA (**Table S1**), again suggesting an underestimation of ManA. Notably, GBV023, GBV034 and ALV118 strains consistently contained two Glc residues, one of which is located in a lateral branch, suggesting that this structural feature remains stable during analysis (**Table S1**). In addition, the corresponding ^1^H and ^13^C-NMR profiles closely matched those previously reported for the EPS produced by strain KYGT207 [7], supporting the conclusion that these four strains synthesized the same EPS structure. Strains GBV030 and GBV036 were assigned to aEPS-I group based on EPS composition profiles similar to that of strain YAS34, with average of 3.0 Glc, 2.2 Gal, and 0.84-1.0 GlcA (**Table S1**). The previously resolved YAS34 EPS structure contains 3 Glc, 3 Gal and 1 GlcA residue per repeating unit (**Figure 1A**). In this structure, one Gal is part of the core chain, whereas two Gal residues are located within a lateral branch - including one pyruvate-substituted residue, which may explain the lower apparent in Gal abundance for GBV030 and GBV036 (**Table S1**). Additionally, GlcA appeared underrepresented in the EPS composition of strain GBV030 (0.84), consistent with the underestimation of ManA observed for ALV104, further suggesting that HPLC-MS analyses tend to underestimate acidic monosaccharides. The primary objective of this study was therefore to validate these EPS groups through the prediction of SOCs involved in aEPSs biosynthesis and comparative analysis of their gene content.

### Prediction of the Supra-operonic Clusters (SOCs)

To predict SOCs involved in EPS biosynthesis in *Rhizobium* genomes, we focused on key genes encoding precursors, primers, decorators and transporters (**Table S2**) and CAZymes (GTs, CEs, GHs and PLs). All CAZymes-encoding genes identified in the genomes of the 87 strains were semi-manually curated by CAZy database experts to ensure high-quality functional annotation (**Table S3**). Additional key genes were annotated based on the presence of Pfam domains [33] as listed in **Table S2**. Key genes were subsequently grouped into SOCs based on their genomic proximity. Starting from any identified key gene, neighboring key genes and separated by fewer than 10 intervening genes were assigned to the same SOC. A scoring system was developed to assess SOC completeness according to functional gene composition. One point was assigned for each transporter gene of the Wzx/Wzy-dependent system (*pssLTNP*), one point for the presence of priming genes (*pssAB*), one point for the presence of at least one GT gene, and 0.5 point for the presence of at least one decorator. The resulting score therefore reflected both the diversity and completeness of the biosynthetic and transport machinery encoded within each predicted SOC, with higher scores indicating more complete EPS-associated systems. Hereafter, all SOCs discussed are referred to as EPS-SOCs.

Across all analyzed genomes, a total of 5,422 SOCs were predicted; however, the vast majority (87%) scored below 2 and lacked transporter genes (**Table S4**). The number of predicted SOCs per genome ranged from 45 to 80 (**Figure 3** and **Figure S1**). *R. alamii* genomes contained an average 76.2 SOCs, with 67.2 scoring below 2 and 9 scoring ≥ 2. In comparison, other *Rhizobium* species, harbored an average of 62.5 SOCs, of which 53.8 scored < 2 and 8.7 scored ≥ 2. *Sinorhizobium* species exhibited a similar distribution with an average of 61.4 SOCs per genome, comprising 52.7 with scores < 2 and 8.7 with scores ≥ 2. Some species, such as *R. rosettiformans*, *R. glycinendophyticum*, *R. ipomoea*, and *R. skierniewicense*, possessed no more than two transporter genes associated with predicted EPS-SOCs. The lack of some transport components required for the canonical Wzx/Wzy-dependent pathway may have several explanations. First, all genomes were annotated using the Pfam HMM library, for which domain detection relies on predefined significance thresholds; consequently, highly divergent domains may remain undetected. Second, some SOCs have undergone partial gene loss or genomic rearrangements leading to transporter gene translocation. For instance, *R. favelukesii* produces EPS using two SOCs acquired via horizontally gene transfer from *S. meliloti*: with one SOC encoding only *pssL*, and the second containing *pssPNT* genes [11]. Finally, some strains may employ alternative transport systems involving isofunctional proteins that fulfill equivalent roles despite lacking detectable sequence homology to canonical Wzx/Wzy components.

**Figure 3:**
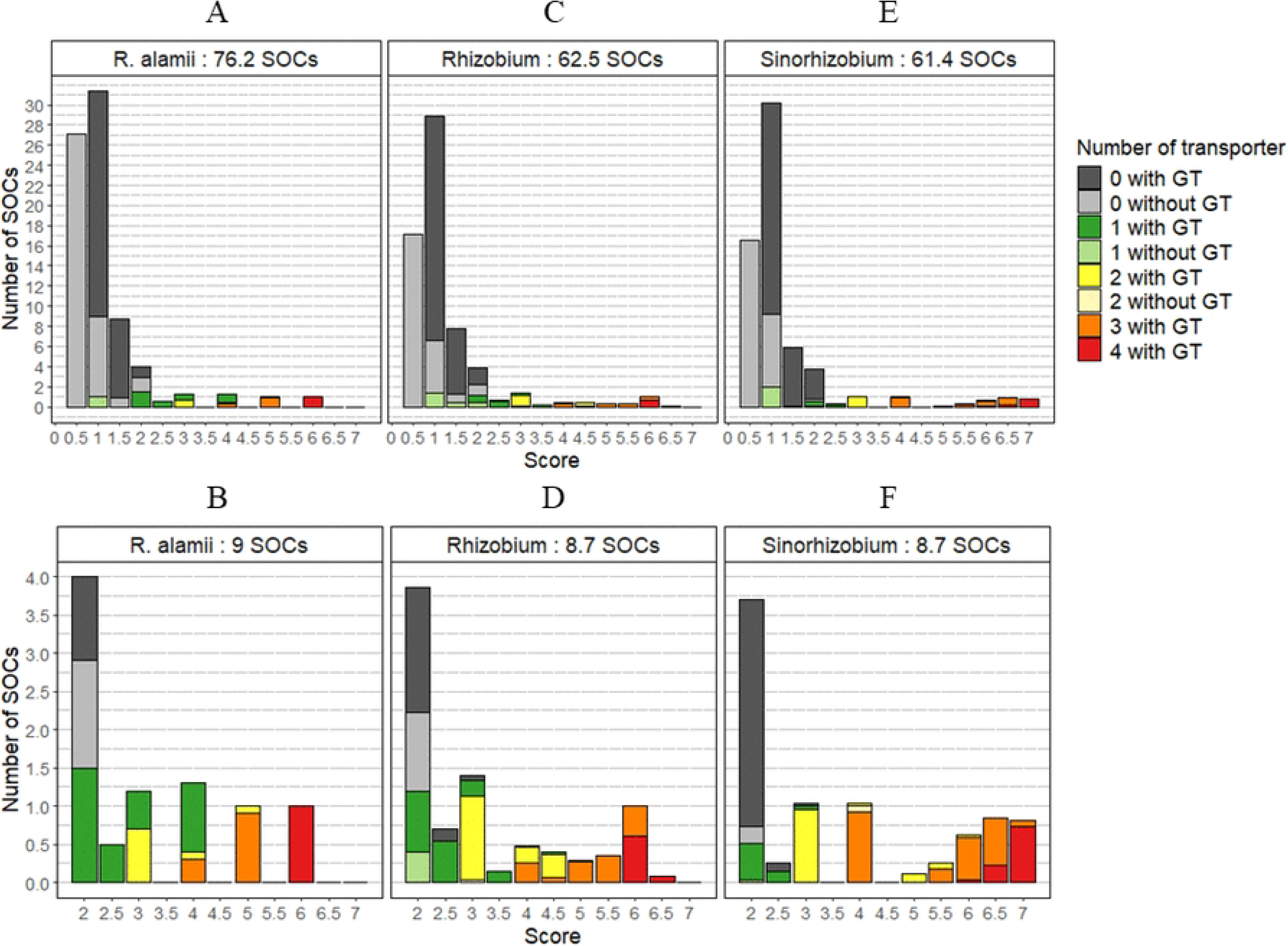
Number of Synthetic Operon Clusters predicted. SOCs are categorized by score (**AB**) across *R. alamii genomes* alone, (**CD**) all *Rhizobium* genomes (excluding *R. alamii*), and (**EF**) *Sinorhizobium* for (**ACE**) all SOCs or (**BDF**) only SOC with a score ≥ 2. Each bar represents the mean number of SOCs with a specific score. Bars are color-coded—grey, green, yellow, orange, or red—based on the number transport proteins from the Wzx/Wzy model (PssLPNT) they contain: 0, 1, 2, 3, or 4, respectively. Bars are filled with solid colors for SOCs containing GTs and with transparent colors for those lacking GT. The numbers next to the genera represent the mean of total SOCs number in the genomes (**ACE**) or only of SOCs with a score ≥ 2 (**BDF**).

To ensure meaningful comparisons of SOCs associated with the Wzx/Wzy-dependent pathway, only SOCs with a predicted score ≥ 2.0 were retained for further analyses. This threshold allowed the inclusion of partial but functionally relevant SOC, such as the translocated repeating unit initiators (*pssA* and *pssB*) in Rlv3841 [10], as well as the second SOC of *R. favelukesii* LPU83, involved in sEPS-I synthesis, which contains only the Wzx transporter and reached a score of 2.5. Following this filtering step, a final dataset of 743 SOCs, was obtained, corresponding to an average of 8.6 retained SOCs per genome.

### Networking predicted and literature-derived SOCs

The 743 SOCs with scores ≥ 2.0 predicted across the 84 genomes analyzed in this study, were systematically compared based on their gene content. Pairwise Jaccard distances were calculated using a modified version of the approach described by Holt et al. [32] SOCs were subsequently represented as similarity networks, in which each node corresponded to a SOC, and edges connected SOC pairs shearing a similarity value (1 - Jaccard distance) above a defined threshold. Similarity networks were generated using thresholds ranging from 0.02 to 0.2. Increasing the similarity threshold progressively fragmented the global network into a larger number of disconnected subnetworks, which were numbered based on their SOC abundance (**Figure S2A**). A threshold of 0.1 was selected to visualize the subnetworks (**Figure 4A**), based on the stability and the clear separation between them (**Figure S2B**). At this threshold, the 743 analyzed SOCs were distributed as follows: 41 remained as singletons, 18 formed isolated pairs, and the remaining 684 clustered into 36 subnetworks. These subnetworks displayed three main organizational patterns: 22 contained exclusively *Rhizobium* SOCs, 6 contained exclusively *Sinorhizobium* SOCs, and 8 contained SOCs from both genera (**Figure 4B**).

**Figure 4:**
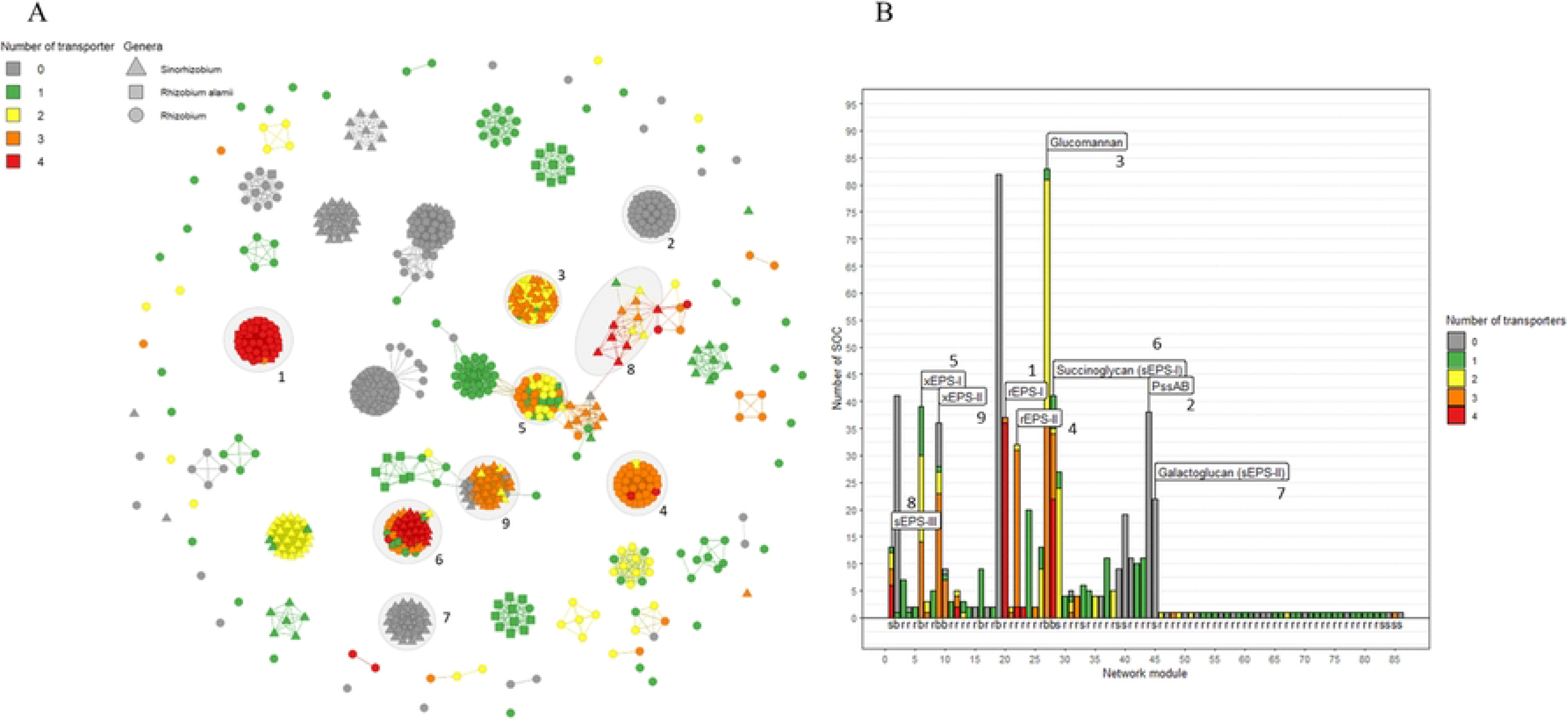
Functional annotation of SOC similarity networks. (**A)** Network of SOCs based on a similarity threshold exceeding 10% between SOCs. Each node corresponds to a SOC identified in the 87 genomes analyzed, with different shapes: triangles for *Sinorhizobium* genomes, circles for *Rhizobium* genomes, and squares for *R. alamii* genomes. Links between nodes represent the similarity between SOCs. Highly connected SOCs form subnetworks, which are visually grouped and indicated by grey circles and ellipses. (**B**) The number of SOCs in each subnetwork, calculated from the network analysis. Nodes and bars are color-coded —grey, green, yellow, orange, or red— based on the number of transport proteins (PssLPNT) they contain: 0, 1, 2, 3, or 4, respectively. Numbers indicate subnetworks containing SOCs with a known function from Rlv3841 and Sme1021. Letters beneath each bar denote the origin of SOCs within a subnetwork: (r) for *Rhizobium* genomes, (s) for *Sinorhizobium* genomes, and (b) for both genera.

Based on the presence of previously characterized SOCs, described in the literature, we could assign a potential EPS structure to seven of the 36 subnetworks. Subnetworks 1 and 2 corresponded to the rEPS-I and *pssAB* SOCs of Rlv3841, respectively, and were exclusive to the *Rhizobium* genus (**Figure 4B**). Subnetworks 3 and 4 contained the SOCs responsible for glucomannan and rEPS-II synthesis in Rlv3841, and were distributed across both genera or restricted to *Rhizobium,* respectively. Subnetworks 6 and 7 corresponded to SOCs associated with sEPS-I and sEPS-II production in Sme1021, with the former present in both genera and the latter restricted to *Sinorhizobium*. Subnetwork 8 included the SOC encoding sEPS-III of Sme1021– an EPS of unresolved structure involved in osmoregulation [34] –and was composed solely of *Sinorhizobium* SOCs. Some *Rhizobium* strains exhibited SOC values similar only to *Sinorhizobium americanum* and were thus excluded from the subnetwork when applying similarity threshold of 0.1. Notably, subnetwork 2, corresponding to the isolated *pssAB* region physically separated from the remainder of the rEPS-I SOC in the genome, and subnetwork 7, which lacked any PssLPNT transporters, exhibited the lowest scores (score = 2). The low score of subnetwork 7 may be attributed to the fact that the corresponding EPS is exported through an ABC transporter-dependent mechanism rather than through the canonical PssLPNT Wzx/Wzy-dependent system [35]. In contract, the highest-scoring SOCs-generally encoding three to four transport proteins-were predominantly found in subnetworks 1 to 9, with the exception of subnetworks 2 and 7 (**Figure 4B**). Interestingly, subnetworks 5 and 9 did not contain any previously characterized SOC, despite being present in both genera, including strains Rlv3841 and Sme1021. The corresponding SOCs spanning genes *pRL90127*-*pRL90170* in Rlv3841 and *SM_b21220*-*SM_b21274* in Sme1021), achieved scores of 6.0 and 6.5, respectively. These SOCs may be involved in the synthesis of previously uncharacterized EPS structures and hereafter designated xEPS-I and xEPS-II, respectively.

For each strain, the 10 highest-scoring SOCs are presented in **Table S5**. Notably, all previously described SOCs from Rlv3841 and Sme1021 were recovered among these top-scoring predictions, with the exception of the isolated *pssAB* region and the galactoglucan biosynthesis SOC, both of which are not associated with the canonical Wzx/Wzy-dependent pathway.

### Grouping species with their predicted EPS profiles

The composition of predicted SOCs demonstrated a strong correlation with the evolutionary trajectories of *Rhizobium* and *Sinorhizobium* species. This correlation was evidenced by comparing the whole-genome distances, estimated using Mash, and SOC distribution profiles (**Figure S3**). Nevertheless, notable exceptions were observed, potentially attributable to lateral gene transfer events.

An alternative representation of the data as a heatmap (**Figure 5**) further revealed that all analyzed genomes, with the exception of *R. skierniewicense*, contained a glucomannan-associated SOC. This widespread distribution suggests vertical inheritance from a common ancestor, together with strong evolutionary conservation of this biosynthetic system. Beyond glucomannan production, the diversity of EPS-associated SOCs segregated the analyzed *Rhizobium* and *Sinorhizobium* species into four major groups. The first group comprised *Sinorhizobium* species together with, *R. favelukesii* and *R. grahamii.* These genomes shared a characteristic combination of SOCs encoding sEPS-I and the uncharacterized EPS xEPS-II, while most also share additional SOCs associated with sEPS-III and sEPS-II biosynthesis. Few exceptions included the absence of the sEPS-II SOC in *S. fredii* and the lack of sEPS-III SOC in *S. medicae*. In contrast the *Rhizobium* representative within this group lacked both sEPS-II -and sEPS-II-associated SOCs. The remaining *Rhizobium* species could be divided into three additional groups. The first included *R. leguminosarum* and *R. alamii* and related species, and was characterized by the presence of four SOCs corresponding to *pssAB*, rEPS-I, rEPS-II and xEPS-I. The main variation within this group was the absence of the rEPS-II SOC in all four *R. phaseoli* strains and in one of the two analyzed *R. etli* strains. The second *Rhizobium* group was characterized almost exclusively by the presence of the glucomannan SOC, with the exception of *R. daejeonense*, which also harbored a *pssAB* SOC. The third group was distinguished by the presence of the sEPS-I-SOC encoding succinoglycan biosynthesis. Among these species, only *R. smilacinae* possessed an additional xEPS-I SOC. Most species belonging to these groups are known to establish symbiotic interaction with plants, through the coordinated expression of Nod-, Nif-, and EPS-encoding genes. These groupings were therefore further investigated to assess the evolutionary relationship between EPS production and plant symbiosis.

**Figure 5:**
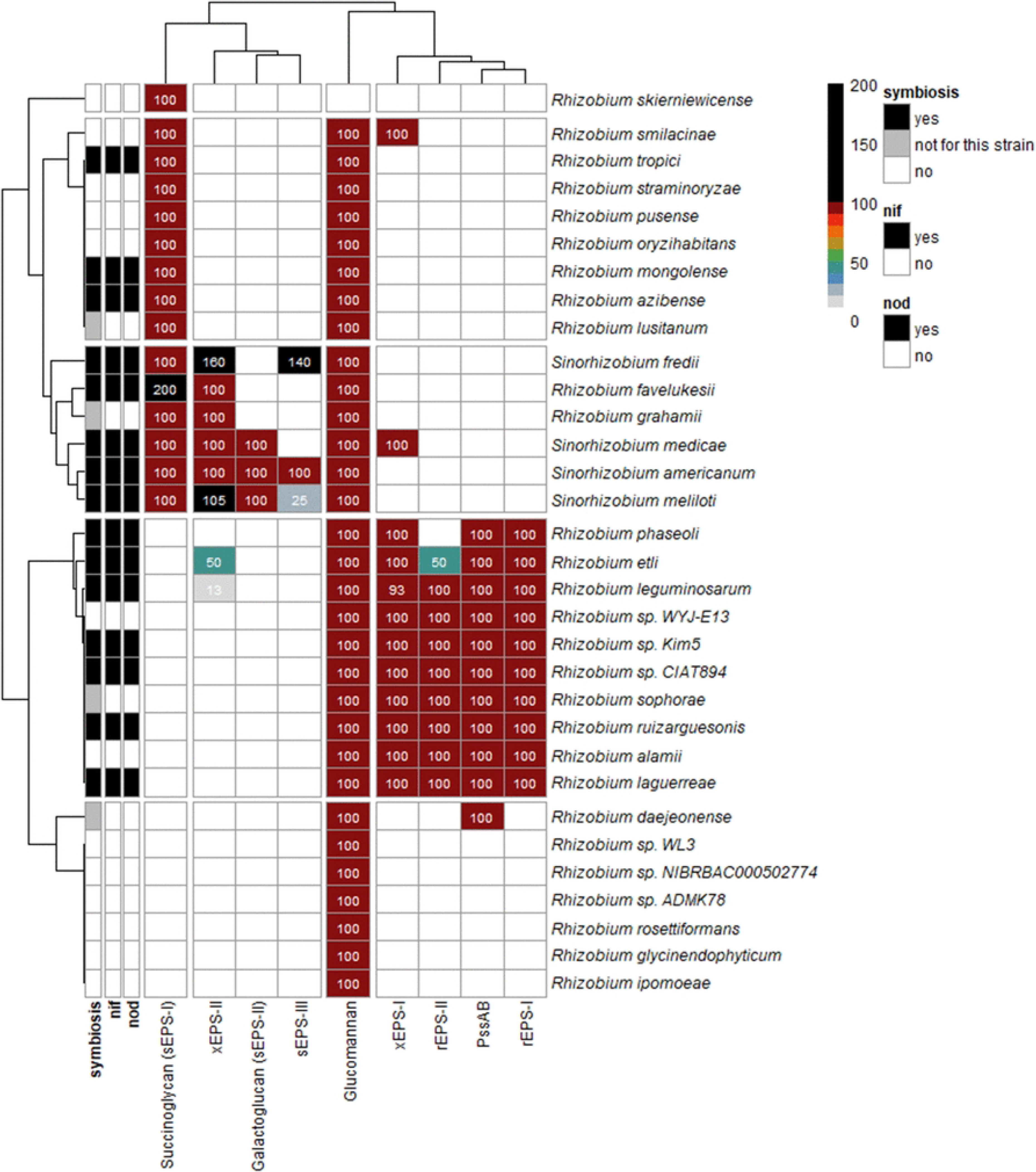
Diversity of SOCs encoding EPS biosynthesis identified in *Rhizobium* and *Sinorhizobium* genomes. Columns represent different EPS-SOC, while rows correspond to each species analyzed in this study. Each color and numerical value indicate the percentage of genomes within a species that contain a SOC in a subnetwork associated with a SOC with a known function. Values below 100% indicate that not all genomes within that species possess a SOC in the subnetwork. Values between 100 and 200% indicate that some genomes within the species contain two SOCs in the subnetwork, and values of 200% indicate that all genomes of that species harbor two SOCs in the subnetwork. Annotations on the left indicate if *nod* genes (*nodABC*) and *nif* genes (*nifHFK*) were annotated in the species genomes in NCBI. The symbiosis annotation indicates if the species were described forming nodulations in the literature. “not for this strain” means that the species was described forming nodulation but not the strain specifically.

### Are plant symbiosis and EPS profiles correlated?

Most *nod* and *nif* genes involved in legume symbiosis within the Rhizobiaceae are located on megaplasmids, also known as symbiotic plasmids (pSym), allowing these traits to be gained, exchanged or lost through horizontal gene transfer [36]. As a result, the capacity for nodulation is not uniformly distributed among all Rhizobiaceae species, with some members lacking the required genes either due to the absence or loss of these plasmid-borne elements. Extensive research has established that rEPS-I is as a key molecular determinant of symbiosis in *R. leguminosarum* and *R. etli*, exerting a major role on interactions with the respective host plants [5]. In contrast, while *R. alamii* and *Rhizobium sp*. WYJ-E13 possess an rEPS-I SOC (**Figure 5**), they lack *nod* and *nif* genes required for the formation of functional nitrogen-fixing root nodules and are therefore considered non-nodulating strains [37]. All strains belonging to the *Sinorhizobium* group exhibited a conserved SOC profile (**Figure 5**) and shared the symbiotic and nitrogen fixation traits characteristic of *S. meliloti* [38,39]. In contrast, *R. rosettiformans*, *R. glycinendophyticum* and *R. ipomoea* possessed only the glucomannan SOC and lacked *nod* genes, consistent with the absence of reported symbiotic relationships with legumes [40–42]. An exception was *R. daejeonense*, for which two strains have been shown to nodulate *Medicago sativa* [43]; interestingly this species was also the only member of this group to possess a *pssAB-*SOC (**Figure 5**). Within the third, which includes *R. tropici*, only half of the 12 species displayed nodulation and nitrogen fixation capabilities [38,44–48]. This heterogeneity in symbiotic potential among species sharing similar SOC profiles indicates that the presence of specific EPS-SOCs alone is not sufficient to predict the capacity of legume nodulation.

### Determinants of rEPS-I structural variation

The *Rhizobium* rEPS-I subnetwork constituted a large dataset, encompassing 37 genomes including the model species *R. leguminosarum* and *R. alamii* (**Figure 6**). Notably, the gene V8V56_0767 (referred as EU184019 in NCBI databases, encoding a GT4) whose deletion impairs EPS synthesis in *R. alamii* YAS34 [31], is located within this SOC. This observation strongly suggests that the rEPS-I SOC of *R. leguminosarum* corresponds to the aEPSs biosynthetic systems characterized in *R. alamii* in the first section of this manuscript (**Figure 1A**). To investigate the genetic determinants underlying EPS structural diversity, we performed a comparative analysis of selected biosynthesis and transport proteins. Variations in EPS structures are expected to depend primarily on (i) PssA, which initiates repeating unit assembly by selecting and transferring the first monosaccharide into the lipid carrier, (ii) GTs, which sequentially elongate the repeating unit, and (iii) decorators, which add substituents such as acetate, pyruvate, or succinate. Proteins involved in EPS polymerization and export are also expected to co-evolve with these biosynthetic components. In particular, PssL is thought to mediate the translocation of properly assembled repeating units, thereby acting as a structural quality-control checkpoint, while PssT facilitates polymerization, with PssN and PssP, which together complete the EPS export machinery.

**Figure 6:**
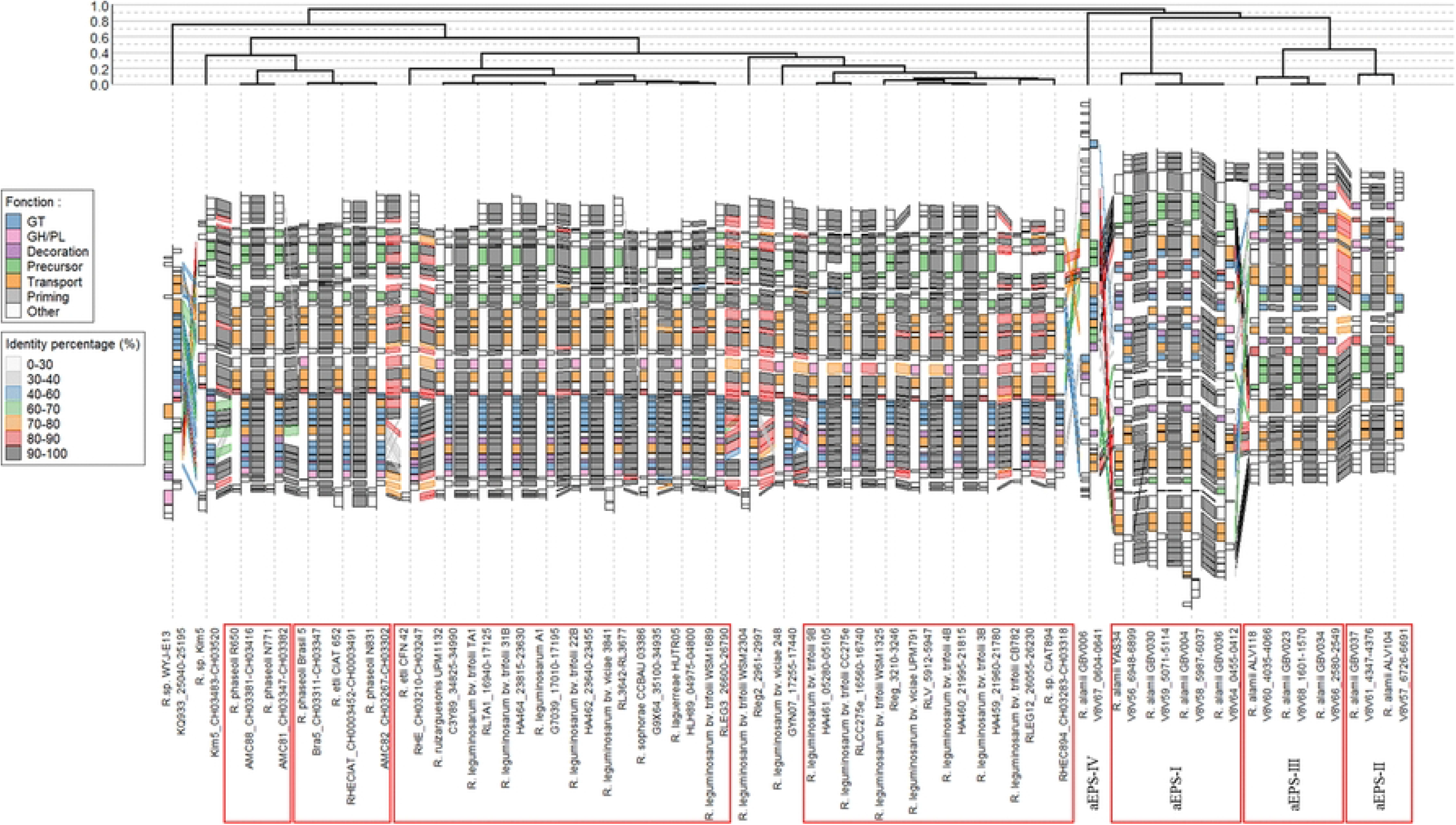
Synteny of 37 SOC within the rEPS-I subnetwork. Each SOC is depicted from bottom to top for each *Rhizobium* and *Sinorhizobium* strain. Individual genes are represented as squares oriented either left or right to indicate their genomic orientation, and are color-coded according to their functional category. Gene homology between SOCs is illustrated by connected squares, which are based on the percentage identity derived from BLASTp analysis. The dendrogram above the synteny plot represents hierarchical clustering on Jaccard distance, calculated using GT, transporter, and decoration genes. Red squares highlight groups of SOCs predicted to produce the same EPS structure, based on Jaccard distance and GT content. For each strain, the locus tags of the first and the last annotated genes are indicated above the strain name.

An interspecies comparison of rEPS-I SOCs was realized through a hierarchical clustering based on composition of glycosyltransferases, decorators, and transporters. To better visualize conserved gene organization and potential evolutionary rearrangements, homologous genes and their positions were highlighted between each adjacent SOCs (**Figure 6**). The analysis revealed that all *R. alamii* SOCs formed a coherent and well-separated cluster, distinct from other *Rhizobium* species. Interestingly, these SOCs further segregated into four subgroups that closely matched the predicted aEPS structural types: (i) strain GBV006 associated with the unique aEPS-IV structure; (ii) strains YAS34, GBV030 and GBV036 producing aEPS-I; (iii) strains ALV118, GBV023, and GBV034 associated with aEPS-III; and (iv) strains ALV104 and GBV037 producing aEPS-II. On the left side of the dendrogram (**Figure 6**), several additional subgroups could be distinguished: (i) the highly divergent *Rhizobium sp*. WYJ-E13; (ii) a cluster including *R. etli* CFN42, *R. leguminosarum* Rlv3841 and TA1 strains with additional *R. leguminosarum* strains as well as *R. laguerreae* and *R. sophorae*, all sharing closely related EPS-SOCs and previously reported similar EPS structures, with some other *R. leguminosarum* strains, as well as *R. laguerreae* and *R. sophorae*; (iii) a second cluster composed of other *R. leguminosarum* strains; and (iv) a group consisting predominantly of *R. phaseoli* strains.

To investigate in the influence of the gain or loss of key genes on EPS structural diversity, comparative analyses were performed using species for which EPS structures were partially or fully characterized (**Figure 7A**). *R. alamii* strains displayed substantial variation in GT contents, consistent with the diversity of their corresponding EPS structures. In each strain, the number of GTs closely matched the number of monosaccharides and glycosidic linkages present in the EPS repeating unit: *pssA* plus the 3, 2, 6 and 4 GTs of the respective aEPS-I to aEPS-IV SOCs accounted for the 4, 3, 7 and 5 monosaccharides and associated glycosidic bonds, observed in these structures (**Figure 7B**). The *pssA* gene, responsible for initiating repeating-unit assembly, was notably distinct from its homologs in *R. leguminosarum,* sharing only from 70 to 80%, sequence identity with those of other *R. alamii* strains and other *Rhizobium* species, **Figure S4**). This divergence is consistent with the initiation of EPS synthesis by Gal in the YAS34, ALV118, ALV104, and GBV006 groups, in contrast to the Glc-initiated EPS structures observed in the other *Rhizobium* species analyzed in **Figure S4**. The additional GT2.11 in ALV118, compared to ALV104, is likely responsible for the additional β(1,3)-linked Glc branched onto the Gal residue, forming a lateral branch (**Figure 7B**). Likewise, the distribution of decorators correlated with EPS modifications: all strains carried the *exoZ* and *pssR* genes, consistent with acetate decoration, while only strains YAS34 and GBV006 possessed the *pssM* gene, consistent with the presence of pyruvate in their EPS.

**Figure 7:**
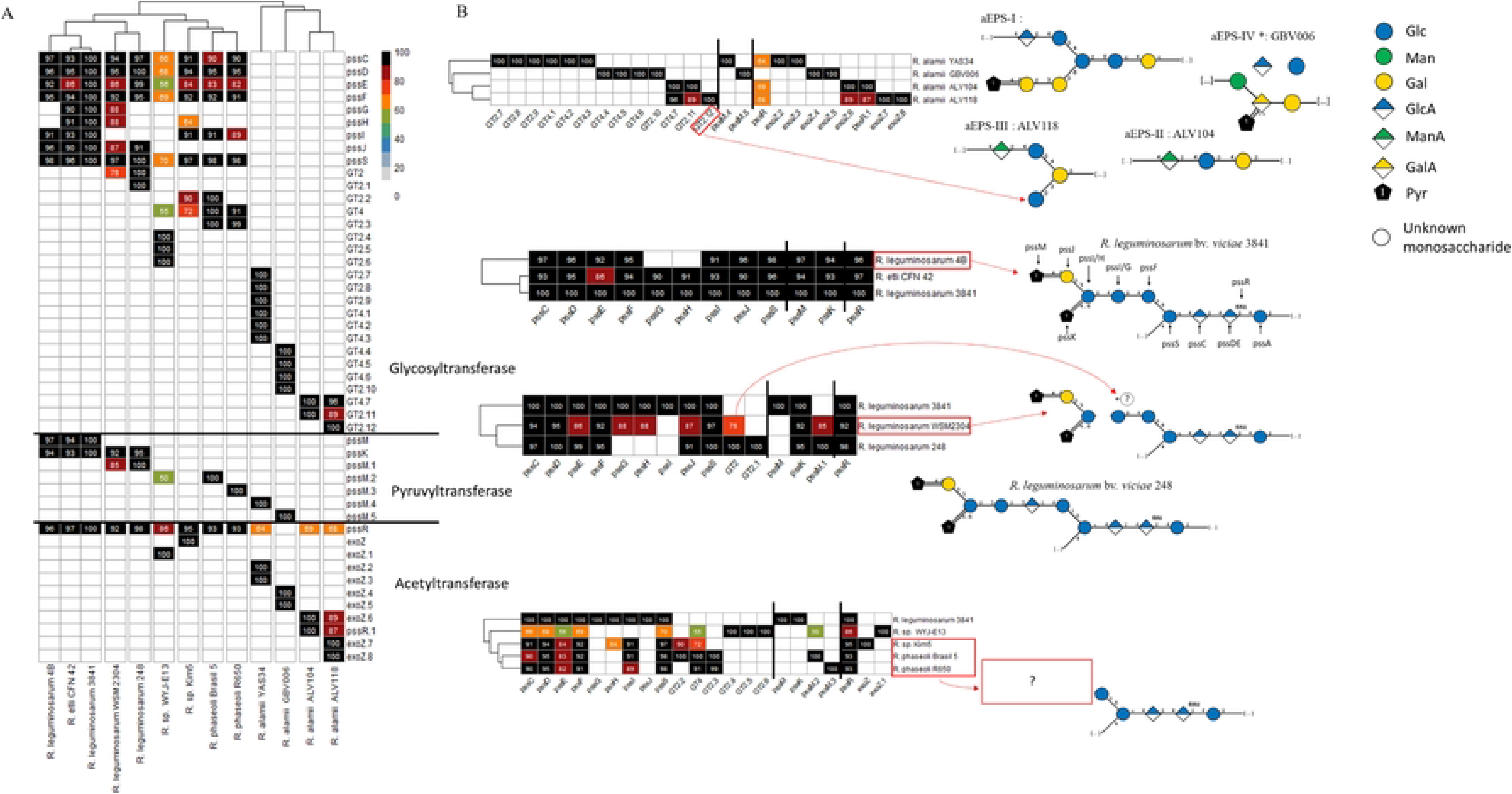
Correlation between key protein content/identity and known EPS structural variations. (**A**) The complete heatmap with a clustering of sequence identity between key proteins: *pss* gene for RLV3841 and each unique GT, pyruvyltransferase and acetyltransferase, represented as rows, and (**B**) Specific subsets highlighting the structural variation (red arrows).

### Correlation between rEPS-I structure and PssL transporter conservation

The *pssLPNT* genes, known from Rlv3841 to encode the core machinery responsible for EPS transport and polymerization, were present in all rEPS-I SOC suggesting that these strains use a conserved export mechanism. However, given the variation in key SOC genes and EPS structures, we hypothesized that evolutionary diversification may also affect the *pssLPNT* genes through structure-associated mutations.

Among these proteins, PssN was the most conserved exhibiting > 80% sequence identity among strains, and clustering into three major groups: (1) *R. alamii* strains, (2) *Rhizobium sp.* WYJ-E13, and (3) the remaining strains (**Figure S5**). This high level of conservation suggests that PssN likely functions independently of EPS structural specificity and is capable of supporting the export of structurally diverse EPS. Similarly, PssP represented the second most conserved component (> 60% identity), displaying a clustering pattern comparable to that of PssN, including the additional duplicated copy identified in *Rhizobium sp.* WYJ-E13 (**Figure S5**). Given that PssP primarily interacts with ATP and PssT during EPS polymerization, it is unlikely to serve as a reliable marker of EPS structural diversity. In contrast, PssT exhibited higher sequence variability with several strains possessing a second copy with less than 20% identity with the primary protein (**Figure S5**). When only the main PssT homologs were compared (**Figure 8A**), *Rhizobium. sp.* WYJ-E13 again formed a distinct lineage, separated from two broader groups corresponding to *R. alamii* strains and the remaining *Rhizobium* species. PssT diversity was markedly higher both within groups (80-100% identities) and between groups (40-60% identities). Further subdivision of these major clusters using a < 90% identity threshold improved the correlation with EPS structural variation. However, some inconsistencies remained between PssT-based clustering and EPS structural variation. For example, strains ALV104 and GBV037 which differed from ALV118, GBV023, and GBV034 strains by the absence of an additional side-branched Glc residue (**Figure 1**) still clustered together. Conversely, certain strains grouped together in the SOC-based analysis of **Figure 6**, like CIAT894, appeared isolated in the PssT-based comparison shown in **Figure 8A** while the previously isolated *Rhizobium sp.* Kim-5 clustered with other strains based on PssT sequence identity.

**Figure 8:**
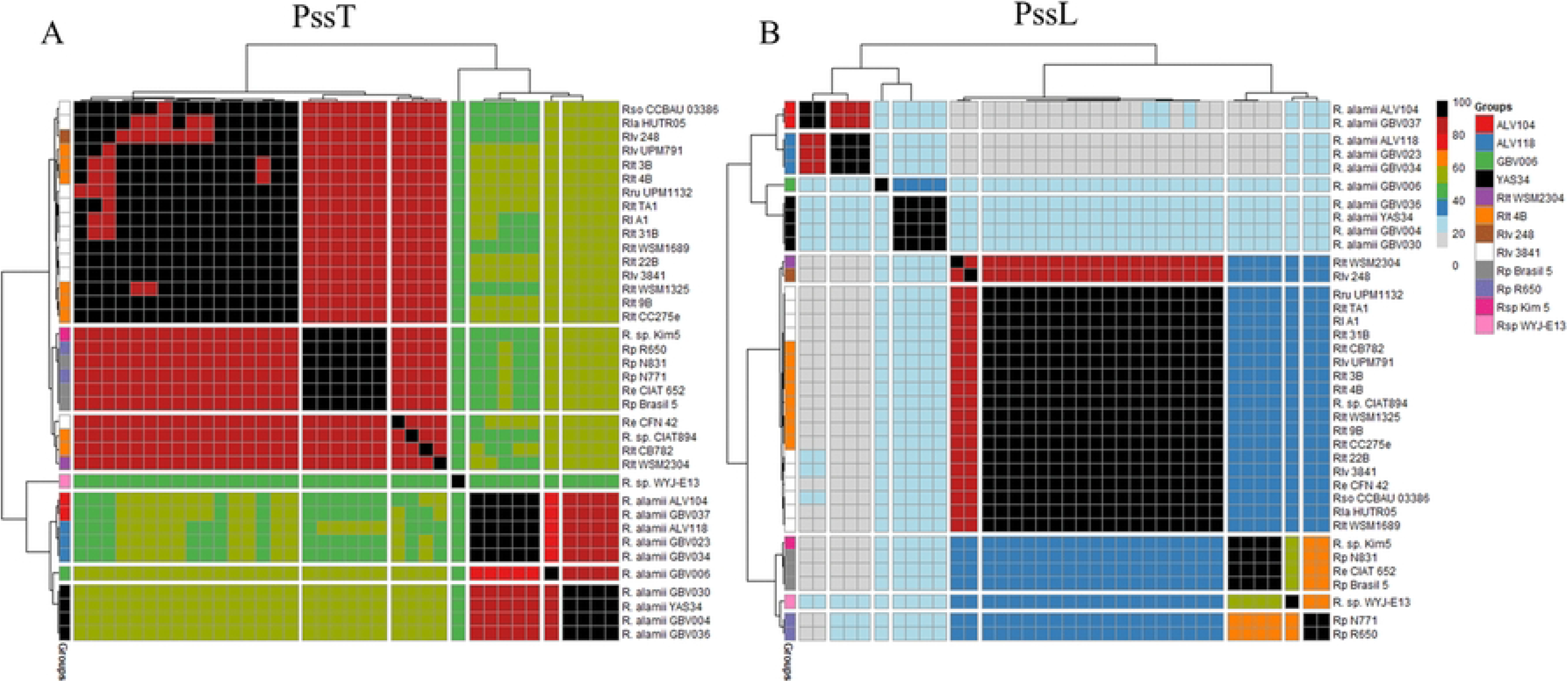
Heatmap of protein sequence identity between SOCs. (**A**) Functional PssT proteins, categorized based on their position within the SOCs and their protein sequence identity relative to Rlv3841 reference protein. (**B**) PssL proteins: colors indicate the percentage identity (ranging from 0 to 100%) between each protein pair, as determined by BLASTp analysis. Row names correspond to the strain from which the proteins were derived. For taxonomic clarity, abbreviations are used: Rlv for *R. leguminosarum* bv *vicieae*, Rlt for *R. leguminosarum* bv. *trifolii*, Rl for *R. leguminosarum*, Re for *R. etli*, Rp for *R. phaseoli*, Rla for *R. laguerrae*, Rru for *R. ruizargenosis*, Rso for *R. sophorea*. Both heatmap rows and columns were sorted based on Bray-Curtis distance and hierarchical clustering using protein sequence identity comparisons derived from BLASTp analysis. The heatmaps were manually divided into k-mers to separate strains based on identity. The colors on the left of each heatmap represent the 12 SOC groups identified in Figure 6. Strains sharing the same color possess highly similar SOCs and are presumed to produce the same EPS structure.

Our comparative analyses revealed a strong correlation between PssL sequence diversity across *Rhizobium* strains and their corresponding EPS composition. Strains producing identical EPS structures exhibited PssL sequence identities of at least 90%, whereas comparisons between distinct EPS groups showed markedly lower conservation, with sequence identities ranging from 10 to 40% (**Figure 8B**). These observations support the hypothesis that PssL may act as a structural checkpoint involved in the selective export of correctly assembled repeating units. Interestingly, the clustering of *Rhizobium sp*. Kim5 with *R. phaesoli* N831 and Brasil5, as well as *R. etli* CIAT652, suggests that their EPS structures could be more similar than expected despite differences in their gene composition (**Figure 6** and **7B**). To determine whether this relationship also applied to other EPS systems, PssL conservation was examined within the sEPS-I. among the 41 analyzed SOCs, 13 lacked a detectable *pssL* homolog (indicated by yellow and green squares in **Figure S6**). The remaining 28 PssL proteins clustered into two groups corresponding to *Sinorhizobium* and *Rhizobium* strains, which shared 79% sequence identity (**Figure S7**). This relatively high level of conservation likely reflects the production of structurally similar succinoglycan in both genera (**Figure 1**). Notably, *S. fredii* SOCs did not contain a canonical *pssL* gene and no gene associated with EPS synthesis in the literature. Comparative analyses between all *S. fredii* SOCs and the Sme1021 sEPS-I SOC, identified a single candidate gene encoding a distant *pssL* homolog (SFHH103_0599) sharing 27% sequence identity (**Figures S8**). Functional inactivation of this gene could determine whether it participates in EPS export in *S. fredii*, which would further suggest that the presence of *pssM* together with two additional GTs may account for the distinct EPS structure observed in this species (**Figure 1**).

These findings underscore two key principles governing EPS biosynthesis. GT content is strongly associated with EPS structural diversity. Second, PssL sequence identity closely correlates with EPS composition among *Rhizobium* strains. Together, these observations support the quality-control model proposed by Ivashina and Ksenzenko [10], in which the transporter machinery acts as a molecular checkpoint ensuring the fidelity of EPS export.

## Conclusion

In this study, we developed a gene-centric framework combining Pfam and CAZy annotations to systematically identify EPS-associated supra-operonic clusters (SOCs) across *Rhizobium* and *Sinorhizobium* genomes. The approach successfully recovered all previously characterized EPS-SOCs and identified two widely distributed, previously undescribed SOC types, designated xEPS-I and xEPS-II. Comparative analyses showed that SOC composition, particularly GT repertoires and Wzx/Wzy-associated transport proteins, strongly correlated with known EPS structural diversity. Notably, PssL sequence conservation closely matched EPS similarity, supporting its proposed role as a structural checkpoint during polysaccharide export.

Although accurate prediction of complete EPS structures remains limited by the broad functional diversity of GT2 and GT4 families, our framework reliably identified candidate EPS biosynthetic systems and enabled clustering of strains producing related EPS. Beyond known systems such as rEPS-I and succinoglycan, the discovery of conserved but uncharacterized SOCs highlights the extent of unexplored EPS diversity within rhizobia.

Overall, this study provides a scalable computational strategy for exploring EPS biosynthetic potential from genomic data and prioritizing candidate systems for experimental characterization. The framework is readily adaptable to other bacterial taxa and offers a foundation for linking EPS biosynthesis, evolution, and ecological function across microbial communities.

## Material and Methods

### Genome selection

The genomes analyzed in this study, excluding the 10 newly sequenced *R. alamii* genomes, were retrieved from GenBank at NCBI. These included 47 genomes from *Rhizobium* species and 27 from *Sinorhizobium* species (**Table S6**). Additionally, we sequenced and submitted 10 *R. alamii* genomes to GenBank under the BioProject PRJNA1082192, corresponding to strains YAS34, ALV104, ALV118, GBV004, GBV006, GBV023, GBV030, GBV034, GBV036, and GBV037. The genomes of YAS34 and ALV104 were sequenced using both Illumina and Nanopore technologies; those of ALV118, GBV030, and GBV037 were obtained with Illumina technology and GBV004, GBV006, GBV023, GBV034, and GBV036 were sequenced using Nanopore technology. Genome assembly results showed that the genomes of YAS34, ALV104, GBV004 GBV006, GBV023, GBV034, and GBV036 were assembled into three or four circular molecules, whereas the genomes of ALV118, GBV030 and GBV037 remained in contig form (**Table S6**). During the study, several *Rhizobium* strains were assigned new taxonomic names, all new taxonomic names have been listed in **Table S6**.

### Exopolysaccharide (EPS) composition and structure

EPS structures of *Rhizobium* and *Sinorhizobium* strains were retrieved from the CSDB bacterial database via taxonomic search [49] or from published literature (**Table S7**). These included: *Rhizobium leguminosarum* bv. *viciae* 3841, *Rhizobium leguminosarum* bv. *viciae* 248, *R. leguminosarum* bv. *trifolii* TA1, *Rhizobium etli* CFN42 and *Rhizobium favelukesii* LPU83), and *Sinorhizobium meliloti* 1021 EPS-I; *S. meliloti* 1021 EPS-II, and *Sinorhizobium fredii* HH103. EPS structures of three *R. alamii* strains were previously characterized: YAS34 [8], ALV104 [6], KYGT207 [7]. NMR H^1^ analysis confirmed that *R. alamii* ALV118 produced the same structure as strain KYGT207. A partial EPS structure for GBV006 was determined by bidimensional NMR H^1^ analysis. The EPS compositions of five additional *R. alamii* strains (GBV023, GBV030, GBV034, GBV036, and GBV037) were characterized in this study. EPS were produced on Tryptic Soy Agar diluted tenfold (TSA/10) supplemented with 20 g.L^-1^ of glucose. After two days of incubation at 30 °C, the EPS were collected from plates by scraping followed by three rounds of centrifugation at 8000 *g* for 15 min at 4 °C. Supernatant was combined, purified with cold NaCl (0.5 M final concentration) and two volumes of cold ultra-pure ethanol under continuous magnetic agitation, washed in ultra-pure ethanol under agitation for 48 h at 4 °C, centrifuged at 8000 *g* for 10 min at 4 °C, washed for 1 h and lyophilized. The purified EPSs were analyzed by HPLC-MS by PCANS platform (CERMAV-CNRS, 38610 Gières, France). Strains sharing the same monosaccharide composition were assumed to possess a similar EPS structure. All known and inferred EPS structures was rendered using the Symbol Nomenclature for Glycan (SNFG) via CSDB/SNFG structure editor [50], facilitating structure visualization and comparison.

### Genome annotation

Genes required for EPS biosynthesis in *Rhizobium* and *Sinorhizobium* strains were identified based on previously published genomic data. In *Rhizobium leguminosarum* bv. *viciae* 3841, these genes are organized within the *pss* Supra-Operonic Cluster (*pss*SOC) [10], while in *Sinorhizobium meliloti* 1021, they are located in the *exo*SOCs [26]. Gene annotation and identification were performed using the Microscope platform, with database selection optimized to detect genes associated with EPS biosynthesis [51]. Functional domains involved in EPS-related processes were identified using Pfam [33], and domain architectures –representing the linear organization and order of individual domains within each gene– were determined from the InterPro database [52]. Domain order information was explicitly considered to distinguish between genes sharing similar compositions but differing in structural arrangement. These domain architectures were then compiled into a curated dataset of EPS-related genes used for SOC prediction (**Table S2**).

For genome annotations, Pfam version 34.0 was used in combination with the HMMsearch tool from HMMER3 suite [53]. For glycosyltransferases, glycoside hydrolases and polysaccharide lyase (GTs, GHs and PLs), the annotation was produced by the experts maintaining the CAZy classification and database [22]. CAZy curators performed semi-manual validations, to ensure accurate classification of borderline homology cases and are listed in **Table S3**.

### SOC prediction

Genes involved in the EPS biosynthesis were grouped into SOCs based on genomic proximity, with neighboring genes assigned to the same SOC if separated by fewer than 10 genes. Each SOC was assigned a score reflecting its functional completeness and gene composition. A GT contributed one point, regardless of copy number, as GT count varies with EPS repeating-unit complexity. Genes encoding the transport components of the Wzx/Wzy model were identified and annotated *<u>pssL</u>* (flippase, Wzx), *pssT* (polymerase, Wzy), *pssP* (co-polymerase, PCP), and *pssN* (outer membrane EPS exporter, OXP), each assigned one point since all four are required for EPS polymerization and export. The priming genes *pssA* and *pssB* received one point each for their role in initiating EPS biosynthesis. Decoration and precursor genes (e.g., those adding acetate or pyruvate) were considered accessory assigned <u>0.5 point</u> each. Auxiliary transporters (e.g., PrsDE from the Type-I secretion system I), GHs, PLs, and modifiers such as EstA were classified as non-essential and excluded from scoring. Only SOCs with a total score above 0 were retained for further analysis. The resulting SOC dataset was compiled into *csv* files containing detailed information for each selected gene including SOC number and score, genomic localization, Pfam and EC-number annotations, and predicted functional assignments.

### SOC comparison

For each genome, protein sequences from genes assigned to SOCs were extracted and compiled in a *FASTA* file. These sequences were compared pairwise across genomes using *blastp* from the BLAST+ software suite [54], applying an e-value threshold < 0.01. All remaining pairwise comparisons were restricted to SOCs with scores >2 to focus on high-confidence clusters.

### SOC network analysis

To quantify the similarity between SOCs, a modified Jaccard index approach, as described by Holt et al. was employed [32]. For each SOC pair (SOCi and SOCj), a complete Jaccard distance—encompassing all genes within the SOCs—was calculated. This distance was derived from two vectors *V*_i_ and *Vj*, whose dimensions corresponded to the assembly of paired genes. Each vector assigned a score of 1 for genes present in SOC_i_, while the identity percentage (ranging from 0 to 1) was used for genes in SOC_j_. Unique genes were scored as 1 for their respective SOC and 0 for the other. Distances between SOCs were computed based on *V*_i_ and *Vj* using the *vegdist* function of in *vegan* package, with the Jaccard distance (*J_ij_*) [55], calculated according to the formula ‘*J* = 2B/(1+B)’, where B is the Bray-Curtis distance. This formula ensures that *J_ij_*values range between 0 (identical) and 1 (maximally dissimilar).

Distances were converted to similarities measures (S_ij_ = 1 - J_ij_) and used to construct a SOC similarity network, retaining edges above a similarity threshold of 0.1. Network construction and analysis were conducted using the *igraph* package, while *visNetwork* was used for interactive visualization [56,57]. Community detection in the similarity network was performed using *igraph*’s *cluster_walktrap*() function, which identifies densely connected subnetworks based on random walks of four steps by default. Subnetwork assignment for each SOC was determined using the *membership*() function, enabling the delineation of discrete SOC subnetwork for subsequent comparative and functional analyses.

### EPS synthesis profiles and evolution

SOCs subnetworks were assigned a function if they contained SOCs of Rlv3841 or Sme1021, known to be involved in the synthesis of a characterized EPS. The functions xEPS-I and xEPS-II were additionally attributed based on the size of the subnetwork and the presence of four necessary transporters. Each SOC within these subnetworks was subsequently assigned the predicted function. The heatmap was generated by selecting only the subnetworks with an assigned function and grouping each genome by species. For each subnetwork with a predicted function, the number of SOCs per species was summed and normalized by the number of genomes in each species group to obtain a percentage, which was then converted into a matrix. This matrix was visualized using *pheatmap()* function of *pheatmap* package [58]. Columns and rows were clustered using the default Euclidian and Bray-Curtis distances, respectively and hierarchical clustering method (complete linkage). For the dendrogram comparison presented in **Figure S3**, a Mash distance was computed on each complete genome using MASH software with a sketch size of 5,000 and k-mer size of 18 [59]. The resulting distance matrix was imported in R, transformed, and subjected to hierarchical clustering using a complete linkage method, followed by conversion into a dendrogram. The subnetwork SOC distance was calculated by counting the presence of every strain in a subnetwork and computing a Bray-Curtis distance with *vegdist()* function. Hierarchical clustering was then performed with *hclust()* function (complete linkage method), and the result was transformed into a dendrogram. Both dendrogram were compared using *tanglegram()* function from the *dendextend* package sorting the branches for both dendrograms [60].

### Synteny comparisons

Within each subnetwork a key Jaccard distances were recalculated by only considering GTs, priming (*pssA*), decoration, transporter (*pssLPNT*) allowing the assembly of a distance matrix. Hierarchical clustering was performed with the *hclust* function from base R package on this Jaccard distance matrix, using the complete method for aggregation. SOC visualizations were produced by an in-house script developed to: (i) represent genes as boxes above or below an axis (according to strand); (ii) compute matching genes between adjacent SOCs and connected their coordinates by colored parallelograms; (iii) optimize SOCs orientation to minimize overlapping crossed parallelograms for improved readability. Visualization scripts were developed using the *ggplot2* and *tydiverse* packages [60, 61], with dendrograms added via the *dendextend* package [63]. To compare key proteins, a heatmap was generated by selecting GTs and decorators within SOCs. Using pairwise *blastp* computed for each protein between SOCs, a data frame of sequence identity was constructed and filtered to retain only pairs with a sequence identity exceeding 50%. This filtered data frame was then used to create a network using the *igraph* package, where each protein was represented as a node and edges were defined by sequence identities above 50%. The *cluster_walktrap()* function, along with membership assignment using default parameters, was used to group proteins into subnetworks based on a minimum identity threshold of 50% of. These subnetworks were used to identify unique protein groups, and a representative protein was selected for each group based on a predefined priority order of strains: Rlv3841, Rlv248, *R. alamii* YAS34, *R. alamii* ALV104, *R. alamii* ALV118, *R. alamii* GBV006, *R. phaseoli* Brasil5, and *Rhizobium sp.* WYJ-E13. Each protein within a subnetwork defined sequence identity relative to the reference protein. A heatmap was subsequently generated using *pheatmap* package based on a matrix comprising each protein from the different strains compared against the reference proteins. For enhance clarity, proteins were manually sorted, and strains were clustered using Euclidian distance and a complete linkage method for aggregation.

### Transporter sequence comparisons

Transporter protein similarities from *blastp* results were displayed as heatmaps using the *pheatmap* package [58]. For PssT proteins, as several strains had multiple *pssT* genes, a manual filtering was performed to keep only the protein with a high identity against the protein of Rlv3841.

## Acknowledgments

JT is grateful to Alain Heyraud f(CERMAV) for EPS biochemical analyses.

## Author contributions

JT, WA and TH conceived and designed the experiments. JT, JL, MLG and NT performed the data analyses. JT, NT, WA and TH wrote the paper.

## Supporting Information

S1 Table. Carbohydrate composition of EPS extracted from *R. alamii* strains.

S2 Table. Genes of interest selected for prediction and their Pfam domain architectures used genomic retrieval.

S3 Table. Curated CAZy annotation made for each genome.

S4 Table. Number of predicted Synthetic Operon Clusters (SOCs).

S5 Table. SOC prediction for each genome.

S6 Table. List of genomes of *Rhizobium* and *Sinorhizobium* strains included in this study.

S7 Table. List of articles for which each EPS structure was retrieved.

S1 Figure. Number and score of SOCs across *Rhizobium* and *Sinorhizobium* species included in this study.

S2 Figure. Modularity and functional annotation of SOC similarity networks.

S3 Figure. Comparison between complete genome of *Rhizobium* and *Sinorhizobium* strains and their known SOC.

S4 Figure. Heatmap of protein identity among PssA and PssB proteins.

S5 Figure. Heatmap of protein identity among PssL, PssP, PssN, and PssT proteins.

S6 Figure. Synteny of 41 SOCs in the sEPS-I (succinoglycan) subnetwork.

S7 Figure. Heatmap of protein identity among PssL proteins in the sEPS-I subnetwork.

S8 Figure. Synteny of sEPS-I SOCs of *S. meliloti 1021*, *S. fredii* HH103 and *R. favelukesii* LPU83.

